# V-type H^+^ ATPase Activity is Required for Embryonic Dorsal-Ventral Symmetry Breaking

**DOI:** 10.1101/2021.10.18.464707

**Authors:** Daphne Schatzberg, Christopher F. Thomas, Patrick Reidy, Sarah E. Hadyniak, Viktoriya Skidanova, Matthew Lawton, Luz Dojer, Shweta Kitchloo, Daniel T. Zuch, Cynthia A. Bradham

**Affiliations:** Department of Biology, Boston University, Boston MA 02215; Molecular Biology, Cell Biology and Biochemistry Program, Boston University, Boston MA 02215; Program in Bioinformatics, Boston University, Boston MA 02215; Biological Design Center, Boston University, Boston MA 02215

**Keywords:** Bioelectricity, p38 MAPK, Nodal, Dorsal-Ventral Axis Specification, Symmetry Breaking, Depolarization, HIF, V-ATPase, sea urchin

## Abstract

The mechanism for embryonic dorsal-ventral (DV) symmetry breaking is idiosyncratic to the species, then converges on polarized expression of BMP signaling ligands. Here, we show that V-ATPase (VHA) activity is an early requirement for DV symmetry breaking in sea urchin embryos. In these basal deuterostomes, DV specification is mediated by ventral Nodal expression that leads to the establishment of a BMP signaling gradient. Nodal expression occurs downstream from p38 MAPK, which is transiently asymmetrically active. We show that VHA activity is required for DV symmetry breaking upstream from both p38 MAPK and Nodal. We rescue VHA-mediated ventralization by enforcing Nodal signaling asymmetry. We identify a VHA-dependent DV voltage gradient and also find that VHA activity is required for hypoxia inducible factor (HIF) activation. However, neither hyperpolarization nor HIF activation account for the dorsalizing effects of VHA, implicating a third unknown pathway that connects VHA activity to p38 MAPK symmetry breaking.

**Graphical Abstract:** 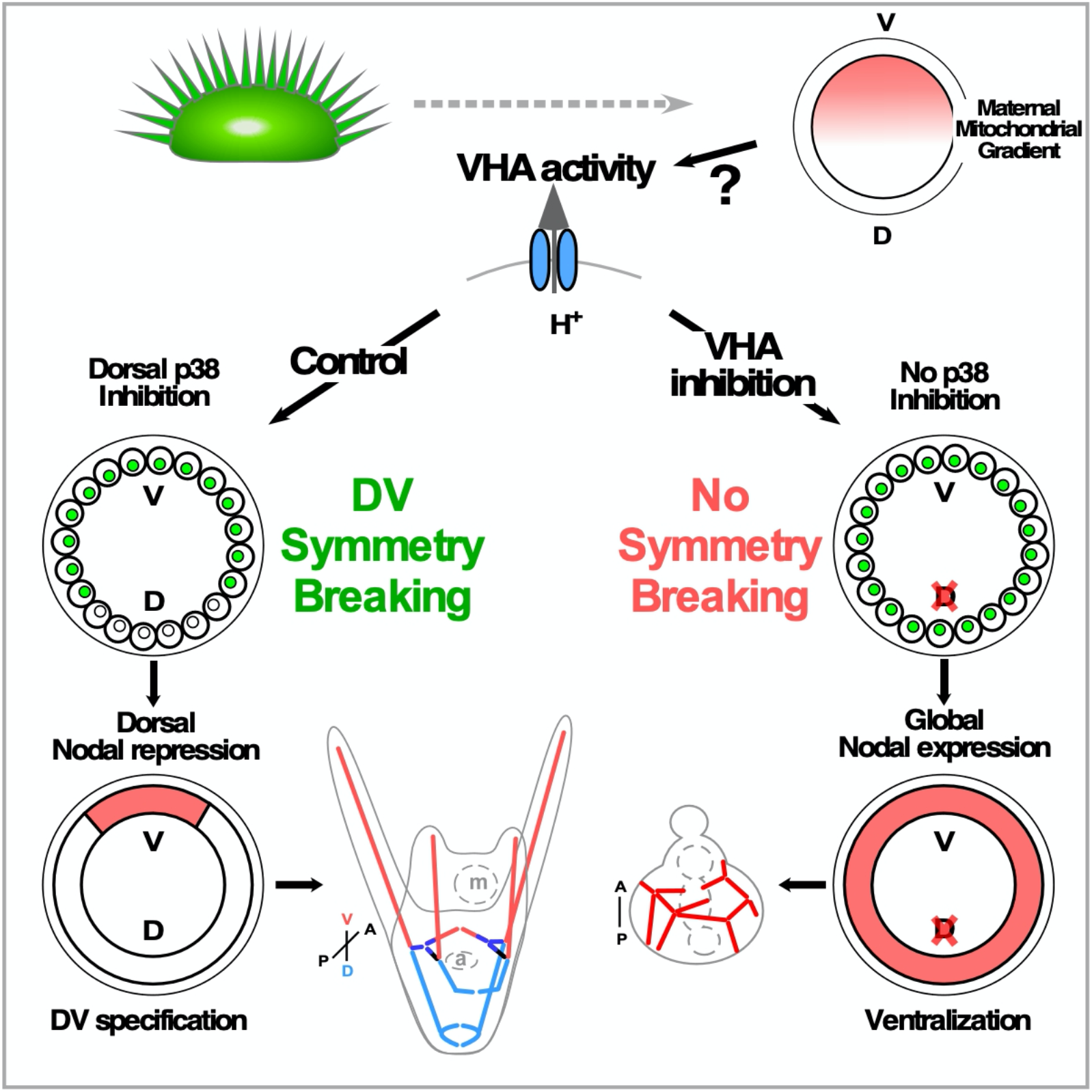

## Introduction

Symmetry breaking in embryogenesis is a fundamental, iterative process that first defines the body axes and germ layers, then subsequently organizes tissues and organs. Polarization occurs at scales ranging from biochemical subunits and polymers through cellular subdomains to the multicellular assemblies that constitute organs and animals. Polarization underlies fundamental processes such as cytoskeletal organization, cell migration, apical-basal and planar cell polarity, vesicular trafficking, stem cell division, neurite formation, and embryonic development; symmetry breaking is thus thought to be the basis for the acquisition of biological complexity (Munro and Bowerman, 2009; Nelson, 2009; Orlando and Guo, 2009; Palmer, 2009; Roth and Lynch, 2009; Tahirovic and Bradke, 2009; Vladar et al., 2009; Wang, 2009; Li and Bowerman, 2010; Yamashita et al., 2010). In developing embryos, axial symmetry breaking occurs by mechanisms that are idiosyncratic to the species and often involve biophysical anisotropies (Munro and Bowerman, 2009; Roth and Lynch, 2009; Houston, 2012; Maitre et al., 2016; Arias et al., 2017; Chen et al., 2018; Zhang and Hiiragi, 2018; Gan and Motegi, 2020; Saadaoui et al., 2020). For the dorsal-ventral (DV) axis, species-specific polarization processes typically converge on BMP signaling gradients that are highly conserved across bilaterians (Little and Mullins, 2006; Bier and De Robertis, 2015).

Sea urchins are basal deuterostomes, and as such, they occupy a key evolutionary position that provides insights into the evolution of chordates and vertebrates and their divergence from radiata and protostomes. As a premier model organism for gene regulatory network analysis that has been studied extensively for over 100 years, sea urchin embryos offer both a breadth of historical investigation and a depth of systems-level and network analyses (Driesch, 1892; Horstadius, 1935; Davidson et al., 2002; Sodergren et al., 2006; Tu et al., 2014; Foster et al., 2020; Hogan et al., 2020; Li et al., 2020; Massri et al., 2021). In sea urchin embryos, DV specification is mediated by zygotic expression of the TGF-β ligand Nodal on one side of the embryo, which defines that side as ventral and leads to BMP gradient formation (Duboc et al., 2004; Bradham and McClay, 2007; Bradham et al., 2009; Lapraz et al., 2009; Su et al., 2009; Saudemont et al., 2010). Nodal is in turn regulated by anisotropically activated p38 MAPK (Bradham and McClay, 2006b; Bradham *et al*., 2009). There is some evidence that the polarized activation of p38 and Nodal is due to asymmetrically distributed mitochondria, whose enrichment spatially coincides with polarized p38 activity in the presumptive ventral region (Coffman and Davidson, 2001; Coffman et al., 2004; Coffman et al., 2009; Modell and Bradham, 2011), although the details of this connection remain unclear.

Here we show that V-type H^+^ ATPase (VHA) activity is required for dorsal-ventral (DV) symmetry breaking in sea urchin embryos. We demonstrate that VHA activity is required for transient p38 inactivation on the dorsal side of the embryo and for the subsequent asymmetric expression of Nodal; when the VHA is inhibited, DV symmetry breaking fails to occur, and embryos are profoundly ventralized. We find that experimentally enforced asymmetric Nodal signaling suffices to rescue the DV axis in VHA-inhibited embryos. We show that while VHA activity is required for a voltage gradient that is normally present along the DV axis, this gradient is surprisingly not required for DV specification. We also find that VHA activity is required for hypoxia inducible factor (HIF) stabilization, but HIF activity is similarly dispensable for DV axis specification. Our results therefore implicate a third, unknown pathway downstream from VHA that regulates symmetry breaking to achieve asymmetric inhibition of p38 MAPK and Nodal to promote dorsal specification.

## Results

### VHA activity is required for normal morphogenesis of sea urchin embryos

The VHA was first isolated from plants and yeast, then classified as a vacuolar proton pump (Lin et al., 1977; Kakinuma et al., 1981; Pedersen and Carafoli, 1987). The VHA is structurally similar to the mitochondrial F_1_F_O_-ATPase but operates in reverse to translocate protons at the expense of ATP (Sun-Wada and Wada, 2015) (Fig. S1A). VHA is a housekeeping gene that is generally ubiquitously expressed (Sun-Wada and Wada, 2015). Each of the VHA subunits was identified in the *Lytechinus variegatus* (Lv) transcriptome (Hogan *et al*., 2020) and found to be expressed at all timepoints with variable expression levels (Fig. S1B; Table S1). FISH analysis confirmed ubiquitous spatial expression of two VHA subunits in the early embryo (Fig. S1C). Bafilomycin A1 (BafA1) and Concanamycin A (ConA) are specific inhibitors of the VHA that bind to the c subunit in the proton-translocating V_O_ domain to prevent the rotation necessary to drive proton movement (Huss et al., 2002; Bowman et al., 2004; Huss and Wieczorek, 2009; Wang et al., 2021). We found that temporal expression levels of the c subunit were unaffected by BafA1 treatment (Fig. S1D).

VHA inhibition with either ConA or BafA1 profoundly disrupted DV axis specification in sea urchin embryos (Fig. 1). VHA-inhibited embryos exhibit strongly radialized skeletons but appear to have gastrulated normally (Fig. 1A). Very similar phenotypes were observed with each inhibitor, suggesting that the radialized phenotype is due to specific inhibition of VHA activity, rather than resulting from an off-target effect. We also combined the two drugs at suboptimal doses and found that they were additive (Fig. S2A), further indicating their overlapping specificity. We attempted to confirm this phenotype by expressing a dominant negative version of *Xenopus* VHA subunit E (Adams et al., 2007), but were unable to identify an effective dose that was not toxic, implying that complete inhibition of the VHA in Lv embryos is incompatible with their survival.

**Figure 1.**
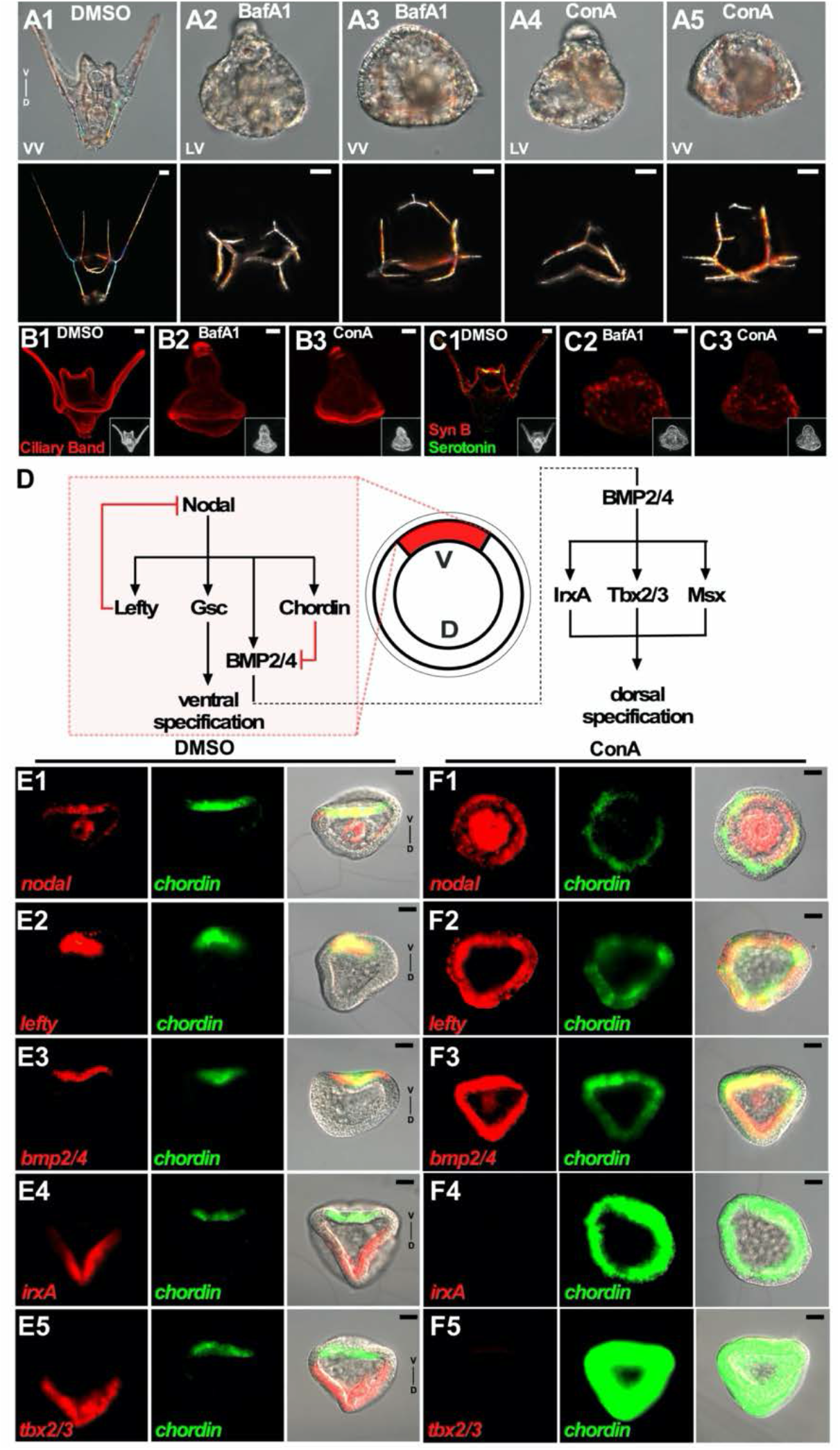
V-type H^+^ ATPase (VHA) activity is required for dorsal specification in sea urchin embryos. **A**. VHA inhibitors radialize the larval skeleton. *Lytechinus variegatus* sea urchin embryos (DIC, upper panels) and their skeletons (birefringence montages, lower panels) are shown at pluteus stage (48 hpf) following treatment with DMSO vehicle (A1) or with the VHA inhibitors Bafilomycin A1 (A2,3) or Concanamycin A (A4, 5) in lateral (LV) or vegetal (VV) views as indicated. **B.-C**. VHA inhibitors ventralize the ectoderm. Larvae are shown at 48 hpf following immunofluorescent labeling for the ciliated band (B) or the serotonin- (green) and synaptotagmin B- (red) positive neurons (C); the corresponding phase contrast images are inset. **D**. A model is shown that depicts the key signals and transcriptional regulators from the ectodermal GRN that specify the sea urchin DV axis. **E.-F**. VHA inhibition ventralizes DV gene expression. Embryos are shown at 18 hpf (late gastrula stage) following fluorescent in situ hybridization (FISH) for the indicated dorsal and ventral genes (see panel D) in control (E) and ConA-treated embryos (F). Right panels show merged signals overlaid on the corresponding DIC images. All scale bars are 20 μm. See also Fig. S1-2.

### VHA activity is required for dorsal specification

To further characterize the effects of VHA inhibition, we assessed ectodermal differentiation markers that exhibit conspicuous differences upon DV perturbation. The ciliary band is a region of apically constricted, ciliated cells that is normally restricted by DV specification to a stripe at the border between dorsal and ventral ectoderm (Fig. 1B1); this band is positioned parallel to synaptotagmin B-positive neurons and their processes, and also encompasses serotonergic neurons in the apical territory (Fig. 1C1). VHA inhibition perturbed both the ciliary band and the neural structures into patterns consistent with ventralization, including the posterior ring of ciliated cells with an apical-most patch (Fig. 1B2-3), diffuse synB neurons, and a complete absence of serotonergic neurons (Fig. 1C2-3) (Duboc *et al*., 2004; Bradham *et al*., 2009; Yaguchi et al., 2010).

DV specification depends on spatially asymmetric Nodal expression, which first directly activates the ventral gene regulatory network (GRN) (Fig. 1D, left), including the Nodal inhibitor Lefty, which is thought to spatially restrict Nodal expression to the ventral territory (Duboc et al., 2008; Muller et al., 2012). Nodal signaling next indirectly activates the dorsal GRN via the induction of both BMP2/4 and Chordin expression; interestingly, BMP2/4 protein relocates from its ventral expression domain to the opposite side, where it signals for dorsal specification (Angerer et al., 2000; Duboc *et al*., 2004; Bradham and McClay, 2006b; Bradham *et al*., 2009; Lapraz *et al*., 2009; van Heijster et al., 2014) (Fig. 1D, right). However, some dorsal genes are detectable when Nodal is inhibited (Bradham and McClay, 2006b; Piacentino et al., 2015; Piacentino et al., 2016), implying that the dorsal network can be at least partially activated in the absence of Nodal expression in Lv embryos, via unknown mechanisms.

VHA-inhibited embryos exhibit spatially unrestricted expression of ventral ectodermal marker genes and an absence of dorsal ectoderm marker genes (Fig. 1E-F), consistent with their ventralized morphology. Importantly, the ventralizing effects of VHA inhibition cannot be explained by a loss of expression for either the Nodal inhibitor Lefty or the dorsal signal BMP2/4 (Fig. 1F2-3). These data confirm that inhibition of the VHA causes complete ventralization of the ectoderm of sea urchin embryos and demonstrate that VHA activity is required for dorsal specification.

### Endocytosis Inhibitors Do Not Perturb DV Specification

VHA is expressed on the plasma membrane of many cell types and thereby regulates the cytoplasmic pH and resting membrane potential of the cell (Gluck, 1992; Narbaitz et al., 1995; Klein et al., 1997; Brown and Breton, 2000; Scarborough, 2000; Nishi and Forgac, 2002; Kawasaki-Nishi et al., 2003; Breton and Brown, 2007; Liao et al., 2007; Pamarthy et al., 2018). However, VHA also acidifies the lumen of membrane-bound organelles such as endosomes, and is required for vesicle acidification, endocytic receptor recycling, and effective membrane trafficking (Nishi and Forgac, 2002; Inoue et al., 2003; Forgac, 2007; Pamarthy *et al*., 2018). To determine whether VHA inhibition promotes ventral specification by interfering with endocytosis, we compared the effects of various endocytosis inhibitors to VHA inhibition (Fig. S2B). 4-bromobenzaldehyde-N-(2,6 dimethylphenyl) semicarbazone (EGA) inhibits late endosome trafficking, while Bromoenol lactone (BEL) inhibits transferrin recycling, a shallow endocytic pathway (de Figueiredo et al., 2001; Gillespie et al., 2013). Neither treatment impacted DV specification as assessed by skeletal morphology (Fig. S2B2-4). EGA treatment resulted in mild stunting of the posterior ventral skeletal elements (ARs, Fig. SB2), while BEL induced only mild skeletal patterning defects at a nontoxic dose (Fig. S2B3-4). These effects do not resemble the phenotypes obtained with VHA inhibition.

Dynasore (DS) blocks dynamin activity, and thereby inhibits clathrin-dependent endocytosis. DS regulates TGF-β signaling in other contexts (Macia et al., 2006; Chen et al., 2009), and was previously described as an inhibitor of DV specification in *S. purpuratus* (Sp) sea urchin embryos (Ertl et al., 2011). While DS treatment had no effect on Lv larval morphology at 6 μM, it had a severe impact at 12 μM, including failure to hatch from the fertilization envelope in greater than 90% of treated embryos. Partially hatched embryos exhibited the most extreme developmental defects, while hatched embryos developed more normally (Fig. S2B5-12). Despite their morphological abnormalities, DS-treated Lv embryos retain an evident DV polarity based on their skeletal morphology (Fig. S2B9-12); critically, these embryos are perturbed but are not radialized and thus do not resemble VHA-inhibited embryos. These results indicate that impaired or inhibited endocytosis does not account for the effects of VHA inhibition, suggesting that the ventralized phenotype observed in VHA-inhibited embryos instead more likely reflects VHA activity at the plasma membrane.

### VHA activity is required for spatially restricted initiation of *nodal* transcription

Since VHA inhibition impacted the spatial expression of Nodal, the initial signal for the DV GRN, we more closely examined the onset of Nodal expression in VHA-inhibited embryos (Fig. 2). VHA inhibition did not affect the timing of Nodal expression, which began at 5.5 hours post-fertilization (hpf), comparable to control embryos. However, Baf1A treatment provoked global onset and maintenance of Nodal transcription, in contrast to the asymmetric expression observed in controls (Fig. 2A-C). At 5.5 hpf in BafA1-treated embryos, global Nodal expression was sometimes enriched on one side (Fig. 2B2), while at subsequent timepoints, Nodal expression in BafA1-treated embryos was global and spatially uniform (Fig. 2B3-4, C2). Thus, VHA activity is not required to regulate the timing of Nodal onset but is required to break symmetry spatially along the DV axis to suppress Nodal expression in the presumptive dorsal territory. Temporal analysis shows that embryos are sensitive to VHA inhibition only until hatched blastula stage (Fig. 2D; Fig. S2C). This profile is strikingly similar to the temporal sensitivity to the DV perturbants NiCl_2_ and SB203580 (SB, a p38 MAPK inhibitor) and corresponds to the established temporal window for DV specification in Lv embryos (Hardin et al., 1992; Bradham and McClay, 2006b). Since p38 MAPK is required to initiate Nodal transcription (Bradham and McClay, 2006b), this temporal profile for VHA inhibition is consistent with VHA inhibition acting before or at the onset of Nodal expression.

**Figure 2.**
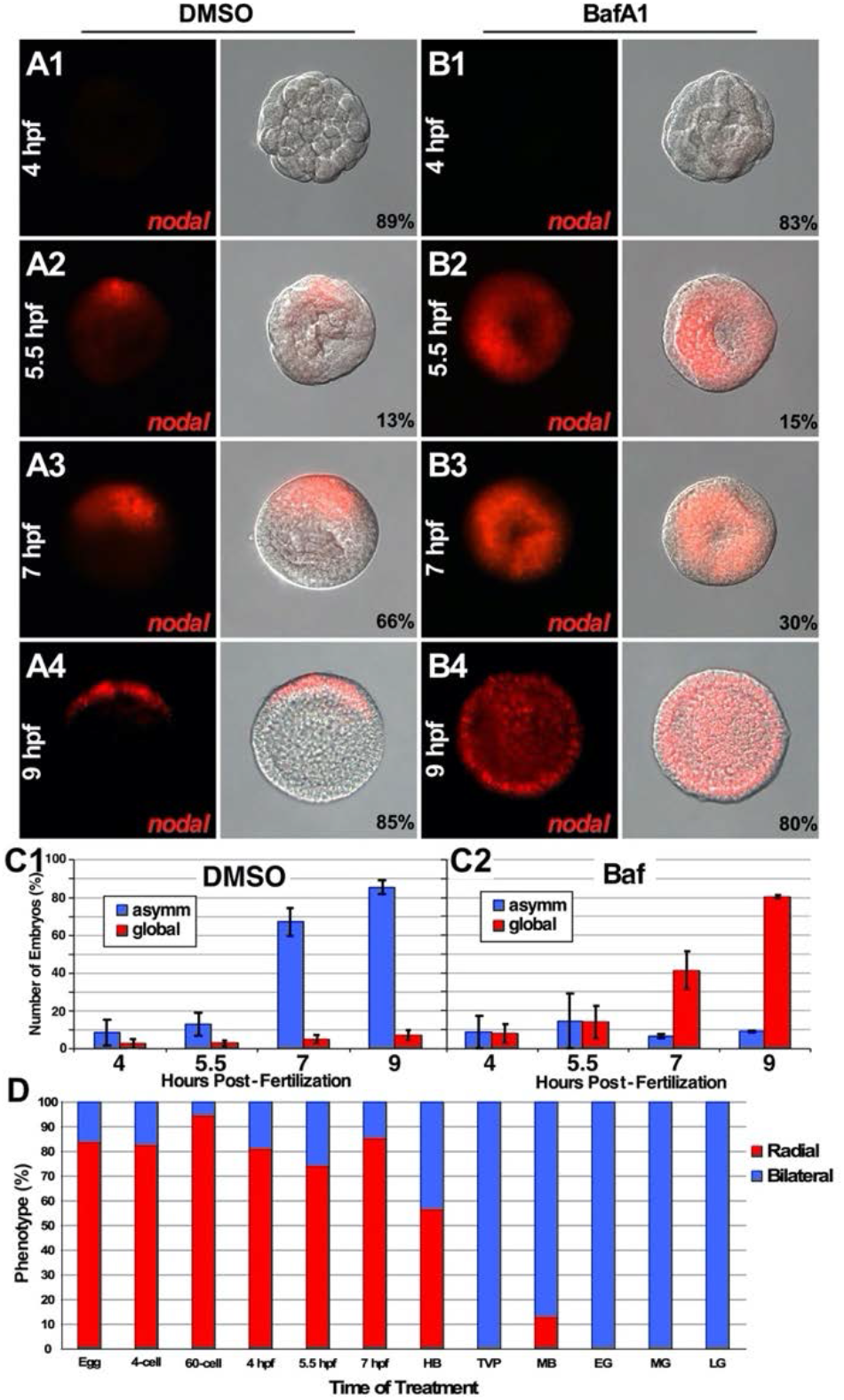
VHA activity is required for spatially restricted Nodal expression. **A.-B**. Nodal onset is correctly timed but spatially unrestricted in BafA1-treated embryos. Embryos treated with DMSO (A) or BafA1 (B) were subjected to FISH for Nodal mRNA at the indicated timepoints. FISH signals alone (left) or overlaid on the corresponding DIC images (right) are shown, with ventral oriented upward when detectable. The percentage of embryos with each expression pattern is shown. **C**. Nodal expression is global in BafA1-treated embryos from the onset. Graphs show the percentage of embryos with asymmetric (blue) or global (red) Nodal expression for DMSO- (C1) or BafA1-treated embryos (C2) at the indicated time points as the average ± s.e.m. from five experiments; n ≥ 45. **D**. Embryos become insensitive to VHA inhibition after hatched blastula (HB) stage. See Fig. S2C for morphologies. Embryos were treated with ConA at the indicated time points, then the fraction of radial (red) and bilateral (blue) morphological outcomes was quantified. HB, hatched blastula; TVP, thickened vegetal plate; MB, mesenchyme blastula; EG, early gastrula; MG mid gastrula; LG, late gastrula. See also Fig. S3.

### Enforced spatial restriction of *nodal* expression is sufficient to rescue dorsal specification in VHA-inhibited embryos

Since VHA inhibition resulted in global Nodal expression, we next asked whether experimentally enforcing spatial restriction of Nodal signaling to one side of the blastula would suffice to rescue DV axis specification in VHA-inhibited embryos. Nodal activates its own expression via a positive feedback loop; since Nodal is secreted, this promotes the spatial spreading of Nodal signaling via a “community effect” (Nam et al., 2007; Range et al., 2007). To experimentally enforce asymmetric Nodal signaling activity, we first inhibited Nodal expression in the zygote via microinjection of a Nodal-specific morpholino (Nodal MO) to prevent positive feedback and thereby block the spatial spread of Nodal signaling. We then expressed a constitutively active form of the Nodal receptor LvAlk4/5/7 Q271D (AlkQD) in one blastomere at the four-cell stage (see Fig. 3D1, red) to confine Nodal signaling to this blastomere and its progeny; VHA inhibitor was applied immediately after the second injection.

**Figure 3.**
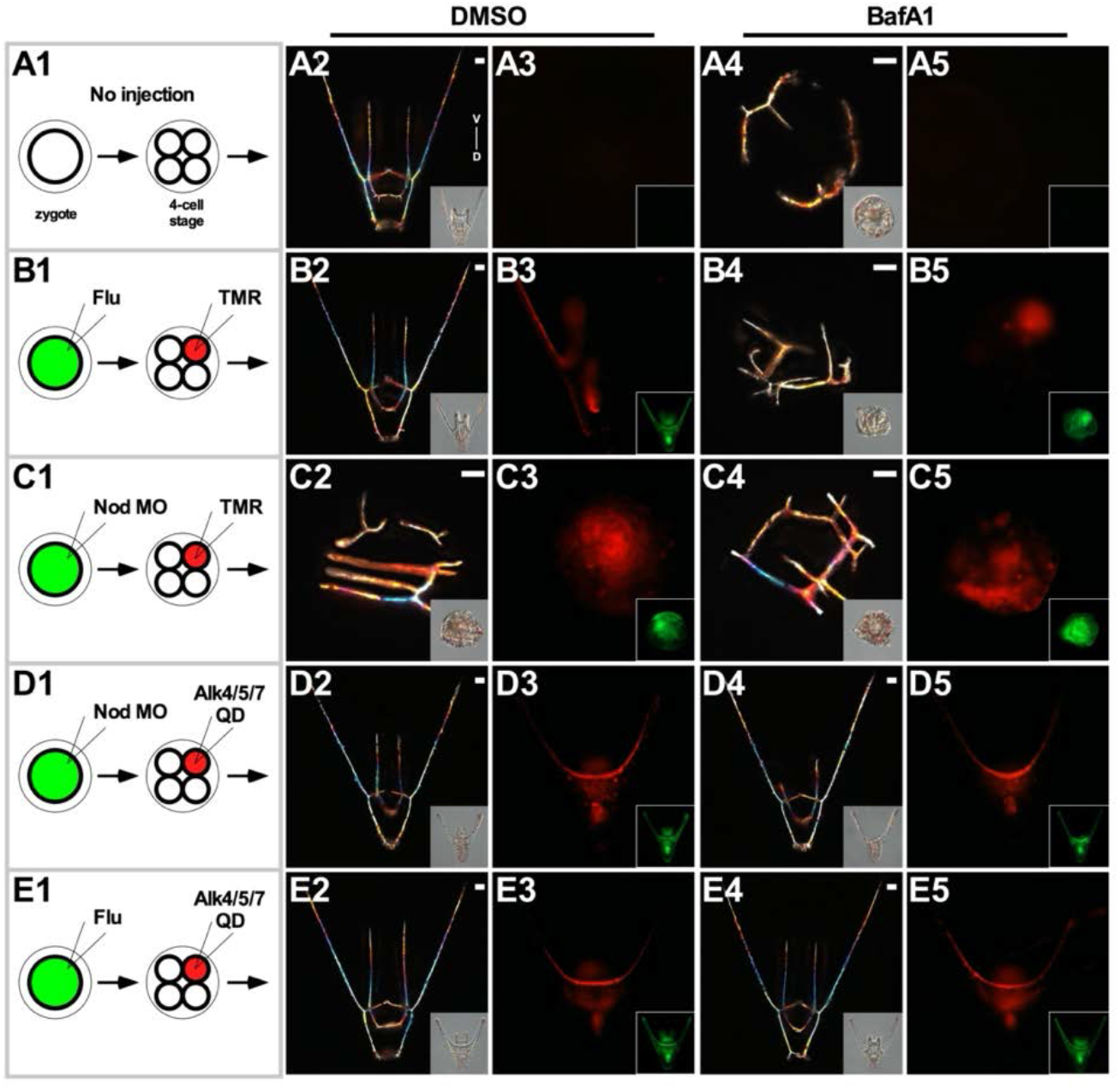
Spatial restriction of Nodal signaling rescues dorsal specification in VHA-inhibited embryos. **A.-E**. Embryos were injected first in the zygote, then in one blastomere at the 4-cell stage, as indicated in the schematics (1), then treated with either DMSO (2,3) or BafA1 (4,5). Control (A), dye-only injected embryos (B), and Nodal MO-injected embryos (C) developed as expected for both DMSO and BafA1 treatments. Asymmetric Nodal signaling (D, E) rescued BafA1-treated embryos. Skeletal birefringence montages are shown at 48 hpf with the corresponding DIC images inset (2,4); for the same embryos, the fluorescence signal from the single injected blastomere (red) is shown with the corresponding signal from the zygotic injection inset (3,5).

Uninjected control and BafA1-treated embryos developed as expected (Fig. 3A). When one blastomere at the 4-cell stage was injected with fluorescent label only, the label was segregated laterally in approximately 70% of control embryos, as expected (McCain and McClay, 1994), without affecting their development (Fig. 3B1-3); label injections also did not impact the BafA1-induced phenotype (Fig. 3B4-5). Injection of Nodal MO alone produced the expected radialized phenotype in controls (Fig. 3C1-3) (Duboc *et al*., 2004; Bradham *et al*., 2009); the combination of Nodal MO with BafA1 treatment also produced radialized skeletons (Fig. 3C4-5). Injection of AlkQD caused the injected blastomere to become specified as ventral in 100% of control embryos, whether the zygote was injected with Nodal MO (Fig. 3D1-3), or with dye only (Fig. 3E1-3); this result agrees with similar experiments in which single blastomeres were injected with Nodal mRNA (Bradham and McClay, 2006b; Bradham *et al*., 2009). Strikingly, AlkQD-injected blastomeres also developed as ventral in BafA1-treated embryos, dramatically rescuing VHA inhibition (Fig. 3D4-5). These results indicate that polarized Nodal signaling is sufficient to rescue DV specification in VHA-inhibited embryos. Importantly, asymmetric Nodal signaling also rescued VHA inhibition without co-injection of Nodal MO (Fig. 3E4-5), demonstrating that asymmetric onset of Nodal expression is sufficient to rescue DV specification when VHA is inhibited. These results indicate that global initiation and not maintenance of Nodal expression is the key functional effect of VHA inhibition, and further corroborate that VHA activity is functionally relevant for DV specification upstream and not downstream from the onset of Nodal expression.

### VHA activity is required for transient p38 inactivation on the dorsal side of the embryo

p38 MAPK activity is required for the onset and maintenance of Nodal expression and for ventral specification (Bradham and McClay, 2006b), which suggested that p38 may be the functional connection between the VHA and Nodal. Interestingly, p38 MAPK is normally globally active in blastula-stage embryos, then is transiently inactivated on the dorsal side of the embryo, just before and during the initiation of ventral Nodal expression (Bradham and McClay, 2006b; Modell and Bradham, 2011), consistent with a model in which p38 MAPK is the critical spatial regulator of Nodal’s expression onset, along with an unknown temporal co-factor (Bradham and McClay, 2006b).

To test for a relationship between the VHA and p38 MAPK, we visualized p38 dynamics in live embryos before and during Nodal expression onset using a GFP-tagged version of Lvp38 MAPK (Bradham and McClay, 2006b). In control embryos, dorsal clearance of p38 occurred between 5 and 5.5 hpf (Fig. 4A), consistent with previous observations (Bradham and McClay, 2006b; Modell and Bradham, 2011); in contrast, p38 clearance did not occur in BafA1-treated embryos (Fig. 4B), indicating that VHA activity is required for dorsal p38 MAPK inactivation. We next performed experiments that combined inhibition of VHA and p38 MAPK. Since p38 inhibition blocked Nodal expression (Fig. 4C-D) and Nodal target gene expression (Fig. 4E) alone or combined with VHA inhibitor, the results indicate that the p38 inhibitor SB203580 operates downstream from the VHA and upstream from the transcription of Nodal to dominantly suppress Nodal expression (Fig. 4C-D) and Nodal-dependent gene expression (Fig. 4E), consistent with a model in which the VHA breaks DV symmetry via dorsal inhibition p38 MAPK.

**Figure 4.**
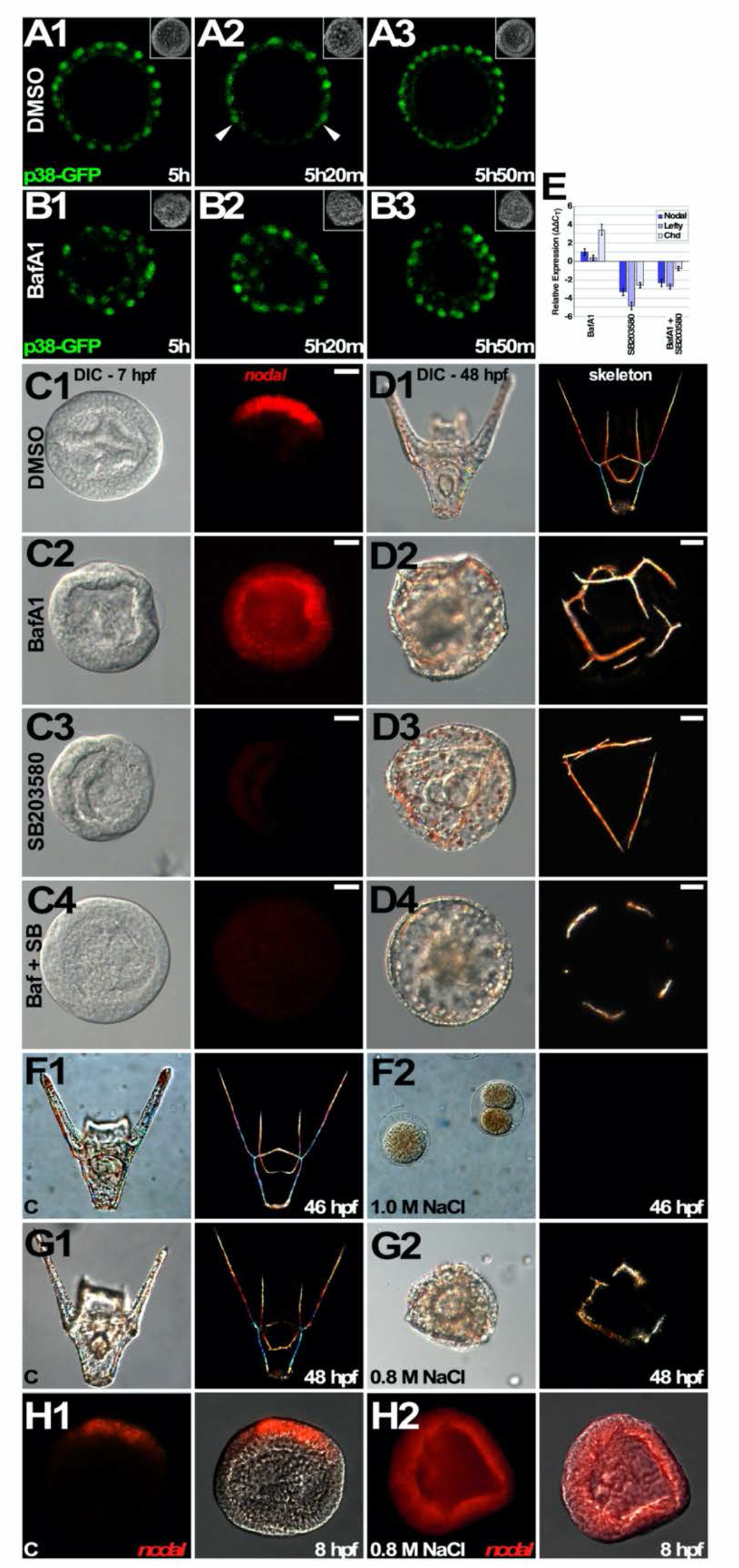
VHA activity is required for transient dorsal p38 MAPK inhibition. **A.-B**. VHA activity is required for dorsal p38 inhibition. Live individual embryos treated with DMSO (A) or BafA1 (B) and expressing p38-EGFP are shown at the indicated timepoints as partial confocal z-stacks, with the corresponding phase contrast images inset. Arrowheads indicate the transient p38 inhibition. **C.-E**. p38 MAPK is functionally downstream from VHA activity. Embryos were treated with VHA inhibitor BafA1 and/or p38 MAPK inhibitor SB203580 as indicated; Nodal expression is shown at 7 hpf (C), and larval morphology and skeleton formation are shown at 48 hpf as DIC and birefringence montage images (D). Expression of the indicated ventral genes was determined by qPCR at 7 hpf in embryos treated with inhibitors alone or together, as indicated (E); the results are shown as the average ΔΔC_T_ compared to DMSO-treated controls ± s.e.m. See also Figure S3. **F**. Embryos treated with 1.0 M NaCl fail to develop past the zygote or 2-cell stage. Control (1-2) and 1.0 M NaCl-treated embryos (3-4) are shown at 46 hpf as DIC (1, 3) and skeletal birefringence montage (2, 4) images. **G.-H**. A minority of embryos treated with 0.8 M NaCl exhibit radialized skeletons (G) and global Nodal expression (H). Control (1-2) and 0.8 M NaCl-treated embryos (3-4) are shown at 48 hpf as DIC and skeletal birefringence montage images (G1-2), and after labeling by FISH for nodal expression at 8 hpf with fluorescence shown alone or overlaid on the corresponding DIC images (H1-2).

### The p38 Inhibitor SB203580 does not inhibit the Nodal receptor

The p38 MAPK inhibitor SB203580 was identified first as an immune modulating drug; efforts to define its target/s led to the discovery of p38 MAPK as its only interactor via affinity chromatography (Lee et al., 1994). SB203580 has been extensively studied and is among the more specific of the kinase inhibitors (Tong et al., 1997; Wang et al., 1998; Davies et al., 2000; Martin-Blanco, 2000; Bradham and McClay, 2006a). Others have raised concerns that in sea urchins, SB203580 does not specifically inhibit p38 MAPK, but instead blocks the Nodal receptor Alk4/5/7, in studies performed in *Paracentrotus lividus* (Pl) Mediterranean sea urchins (Molina et al., 2017). To address this concern, we first tested whether SB203580 inhibits Alk4/5/7 by comparing it with the established Alk4/5/7 inhibitor SB431542 (Inman et al., 2002; Laping et al., 2002) in combination with Nodal overexpression (Fig. S3A). Overexpression of Nodal was sufficient to rescue ventral gene expression in combination with SB203580 but not with SB431542, as measured spatially via FISH in Lv (Fig. S3B), and in a statistically significant quantitative manner in both Lv and Pl embryos (Fig. S3C1-2). This matches the predicted effects, specifically that p38 MAPK is required to induce Nodal expression, while Alk4/5/7 receptor activity functions downstream from Nodal expression (Fig. S3A). Together, these results demonstrate that 20 μM SB203580 does not inhibit Alk4/5/7 in either Lv and Pl sea urchins, and that SB203580 inhibition acts upstream from Nodal, while SB431542 distinctly does not, in both species.

### Hyperosmolarity is sufficient to globally activate Nodal expression

Molina et al. employed osmotic stress as a means of activating p38 MAPK (Molina *et al*., 2017). We similarly tested 1.0 M NaCl exposure and found that this treatment was rapidly toxic to Lv embryos, halting their development at the zygote or 2-cell stage (Fig. 4F). In our hands, 0.8 M NaCl was also largely toxic, but we observed a small number of surviving embryos that exhibited radial morphology and global Nodal expression (Fig. 4G-H), consistent with the interpretation that hyperosmolarity suffices to activate p38 MAPK which in turn suffices to globally activate Nodal expression.

### p38 MAPK inhibition in Pl embryos results in radialized embryos

Molina et al. reported that SB203580 induced unexpected and bizarre phenotypes in Pl, e.g., multiple blastopores and guts, unlike the radialized phenotypes obtained with the Alk4/5/7 inhibitor SB431542, and this was interpreted as reflecting a species difference or an off-target effect (Molina *et al*., 2017). We therefore compared the larval phenotypes of Pl embryos treated with SB203580 and SB431542 and found that each drug similarly perturbs the larva, blocking both DV specification and skeleton formation (Fig. S3D). The effects obtained with SB203580 in Pl embryos are very similar to the effects of SB431542 in Pl (Fig. S3D), and to the previously reported phenotypes in Lv (Bradham and McClay, 2006b), and do not agree with the previously reported, highly unusual effects of SB203580 in Pl (Molina *et al*., 2017).

Together, our results demonstrate that SB203580 acts upstream from Nodal expression and does not inhibit Alk4/5/7 in either Pl or Lv embryos; our hyperosmolarity experiments further support our model that positions p38 MAPK upstream from the onset of Nodal expression. Our findings are not consistent with the reports of cross-reactivity between SB203580 and Alk4/57 (Molina *et al*., 2017); the reasons for these discrepancies remain unclear.

### An endogenous V_mem_ gradient is present across the dorsal-ventral axis and requires VHA activity

Because the VHA translocates protons, it is an electrogenic pump (Gluck, 1992; Brown and Breton, 2000; Scarborough, 2000; Kawasaki-Nishi *et al*., 2003; Liao *et al*., 2007). We therefore visualized the plasma membrane voltage (V_mem_) using the fluorescent reporter DiSBAC (Epps et al., 1994; Adams and Levin, 2006; 2012a; b; Rodriguez-Sastre et al., 2019). We also visualized active mitochondria, which are enriched in the presumptive ventral region of the zygote and embryo (Coffman *et al*., 2004; Modell and Bradham, 2011), to determine the spatial orientation of the DV axis in live blastula-stage embryos. We did not detect differences in mitochondrial ventral enrichment when comparing control and BafA1-treated embryos (Fig. 5A, 5C3). This result was unsurprising since the mitochondrial asymmetry is present prior to fertilization, then maintained by cell division (Coffman *et al*., 2004; Modell and Bradham, 2011). Control embryos exhibit a gradient of plasma membrane depolarization along the dorsal-ventral axis in control embryos between 5 and 6 hpf, such that the ventral ectoderm is relatively depolarized, while the dorsal ectoderm is relatively hyperpolarized (Fig. 5B1-2). In contrast, VHA-inhibited embryos exhibit a flattened V_mem_ gradient, with less dorsal hyperpolarization (Fig. 5B3-4).

**Figure 5.**
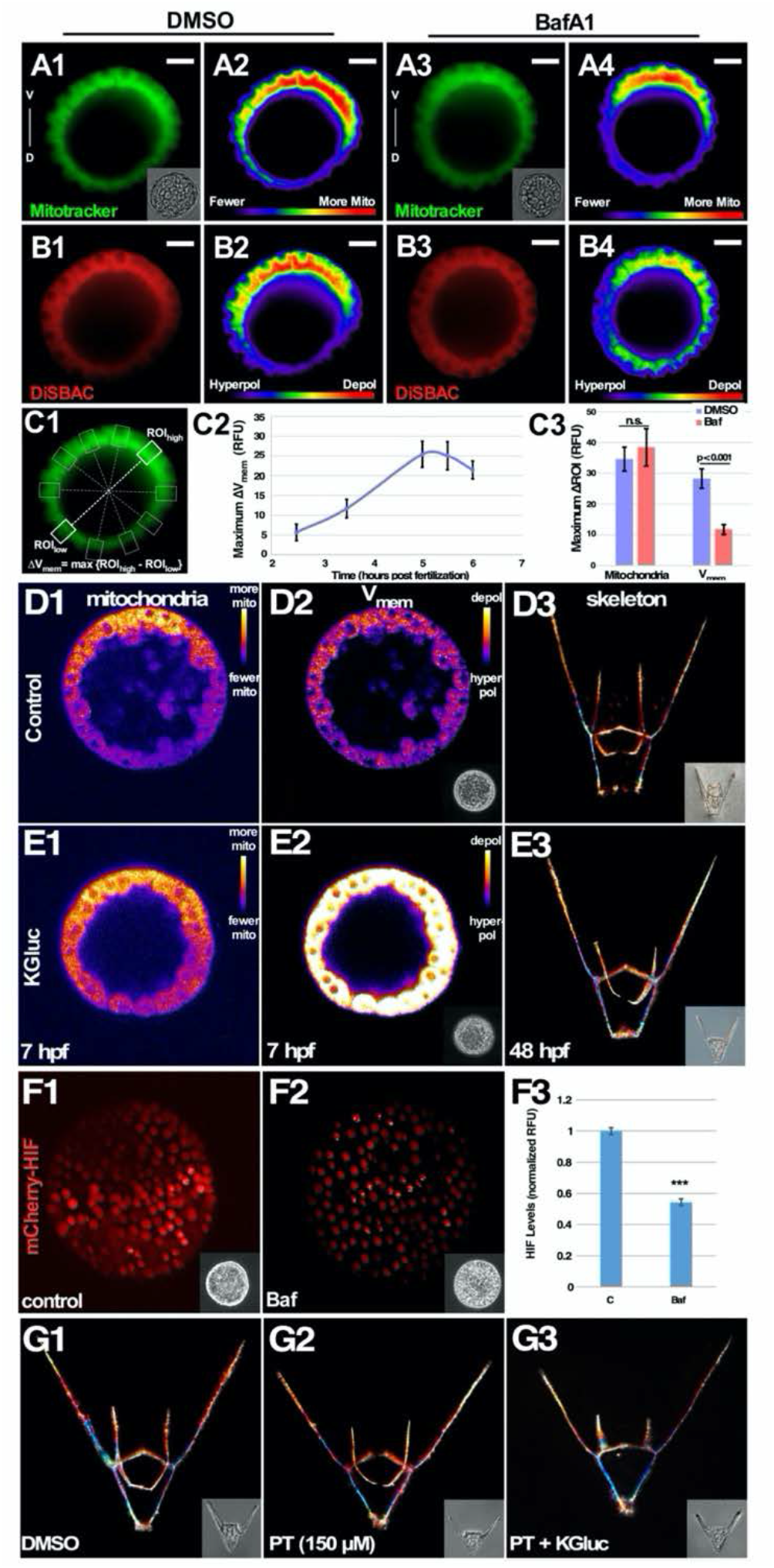
VHA-dependent dorsal hyperpolarization and HIF activation are dispensable for dorsal specification. **A.-B**. VHA activity is required for normal dorsal hyperpolarization. Active mitochondria labeled with MitoTracker Green to mark the DV axis (A) and V_mem_ detected with DiSBAC (B) were visualized together in live blastula-stage embryos treated with DMSO (1-2) or BafA1 (3-4). Fluorescent (1, 3) and corresponding pseudo-colored (2, 4) images are shown. **C**. The gradient of signal was measured as the maximum cross-embryo signal difference, which was quantified by comparing regions of interest (ROIs) on opposite sides of the embryo as indicated (C1). The V_mem_ DV gradient time course (C2) is shown as the average maximum cross-embryo difference along the DV axis ± s.e.m. The mitochondrial and V_mem_ gradients are similarly shown in DMSO- and BafA1-treated embryos at 6 hpf (C3); BafA1-treated embryos exhibited an average of only an 11% maximum difference across the embryo, compared to 28% in controls. p values were determined with paired student t-tests; n.s. not significant. **D.-E**. Hyperpolarization is not required for dorsal specification. Mitochondria and V_mem_ were imaged as in panels A and B in control (D1-2) and potassium gluconate (KGluc, 30 mM)-treated embryos (E1-2) at 7 hpf; the resulting morphology at 48 hpf is shown as skeletal birefringence montages with the corresponding DIC images inset (D3, E3). **F**. VHA activity is required for HIF stabilization. Live control (F1) and BafA1-treated embryos (F2) expressing Sp-mCherry-HIF are shown at 5.5 hpf, with corresponding phase images inset. The quantified results are shown as the average net RFU/embryo ± s.e.m. *** p value < 1×10^−9^ (paired student t-test). **G**. Hyperpolarization and HIF activity, alone or together, are not required for dorsal specification. Embryos were treated with DMSO (G1), the HIF inhibitor PT2385 (PT) alone (G2), or PT together with KGluc (G3), and the resulting morphology is shown at 48 hpf as in panels D3 and E3. See also Fig. S5.

We quantified the maximum difference in fluorescence across the embryo (Fig. 5C1) and found that the V_mem_ gradient arises prior to the onset of Nodal expression at 5.5 hpf (Fig. 5C2) and is significantly depressed by VHA inhibition (Fig. 5C3). The onset of the V_mem_ gradient at 5 hpf coincides temporally with increased expression of VHA subunits F, H, and c (Fig. S1B), possibly explaining the timing of onset for the V_mem_ gradient. These results show that VHA activity is required for hyperpolarization of the presumptive dorsal region of the blastula-stage embryo.

To test whether the p38-to-Nodal pathway regulates the V_mem_ gradient, we measured membrane voltage in embryos in which either p38 MAPK, Nodal, or the Nodal receptor Alk4/5/7 was inhibited. While each treatment resulted in radialized embryos as expected, none exhibited a significantly different V_mem_ gradient when compared to controls (Fig. S4G), demonstrating that the endogenous voltage gradient does not require p38 MAPK activity, Nodal expression, or Nodal signaling, and therefore is not downstream from this pathway.

### Dorsal specification does not require hyperpolarization

Our results suggested that dorsal hyperpolarization may negatively regulate p38 MAPK to break DV symmetry. Since the presumptive dorsal side is relatively hyperpolarized in a VHA-dependent manner at the time of p38 activity asymmetry and Nodal expression onset, we tested whether blocking that hyperpolarization would block dorsal specification. We used potassium gluconate (KGluc) treatment to depolarize the embryo. External K^+^ exposure imbalances the internal positive charge and depolarizes cells (Adams and Levin, 2006; 2013). However, despite strongly depolarizing the embryo (Fig. 5D2, E2), this treatment did not block DV axis specification, and only mild skeletal patterning defects were produced (Fig. 5D3, E3). The results show clearly that depolarization does not suffice to inhibit symmetry breaking or dorsal specification; we thus conclude that hyperpolarization is not required for and therefore cannot account for VHA-mediated dorsal specification.

### HIF contributes only mildly to dorsal specification

Hypoxia inducible factor (HIF) is a redox-sensitive bHLH-PAS transcriptional regulator that is required for mammalian embryonic development (Lee et al., 2019; Corrado and Fontana, 2020). HIF is stabilized under hypoxic conditions by a lack of hydroxylation, and is otherwise degraded under normoxia (Fig. S5A) (Martinez-Saez et al., 2017; Ban et al., 2021). Previous studies in sea urchins showed that SpHIF is dorsally stabilized; however, while HIF plays a supportive role in dorsal specification, it alone is not sufficient to specify dorsal (Chang et al., 2017). VHA is a HIF activator in other contexts (Miles et al., 2017); we therefore tested whether VHA inhibition blocks HIF stabilization using an SpHIF-mCherry fusion protein (Chang *et al*., 2017). We found that dorsal HIF activation indeed requires V-ATPase activity (Fig. 5F). This observation suggested that HIF and hyperpolarization may additively drive dorsal specification downstream from the VHA.

We tested this hypothesis by blocking HIF stabilization with the drug PT2385 (PT), which interferes with HIF dimerization and transactivation (Fig. S5A) (Martinez-Saez *et al*., 2017; Ban *et al*., 2021). Consistent with prior functional studies in Sp sea urchins (Chang *et al*., 2017), we found that HIF inhibition had no discernable impact on DV specification in Lv embryos (Fig. 5G2). However, PT was sufficient to complement a suboptimal dose of BafA1 to radialize the skeleton, showing that HIF inhibition can synergize with weak V-ATPase inhibition (Fig. S5B) and therefore contributes to dorsal specification, albeit weakly. We found that embryos treated with combined PT and KGluc also had normal DV specification and only mild skeletal patterning defects (Fig. 5G3). These results show that alone or together, loss of hyperpolarization and/or HIF inhibition does not suffice to inhibit dorsal specification; thus, these two downstream effects of the VHA cannot account for its DV symmetry breaking effects.

## Discussion

Embryonic dorsal-ventral symmetry breaking is a crucial event for all bilaterian animals that is achieved by different mechanisms, depending on the species, that converge on polarized BMP signaling. Sea urchin DV patterning is currently understood to be controlled by a combination of maternally inherited factors, such as the maternal mitochondrial asymmetry, and subsequent embryonic signaling events that lead to the asymmetric onset of Nodal expression, which in turn activates expression of BMP2/4 and the BMP inhibitor Chordin to drive DV specification. The results herein are consistent with a model (Fig. 6) in which VHA activity, possibly downstream from the maternal mitochondrial asymmetry, mediates dorsal p38 MAPK inhibition to break symmetry along the DV axis upstream from Nodal expression in sea urchin embryos.

**Figure 6.**
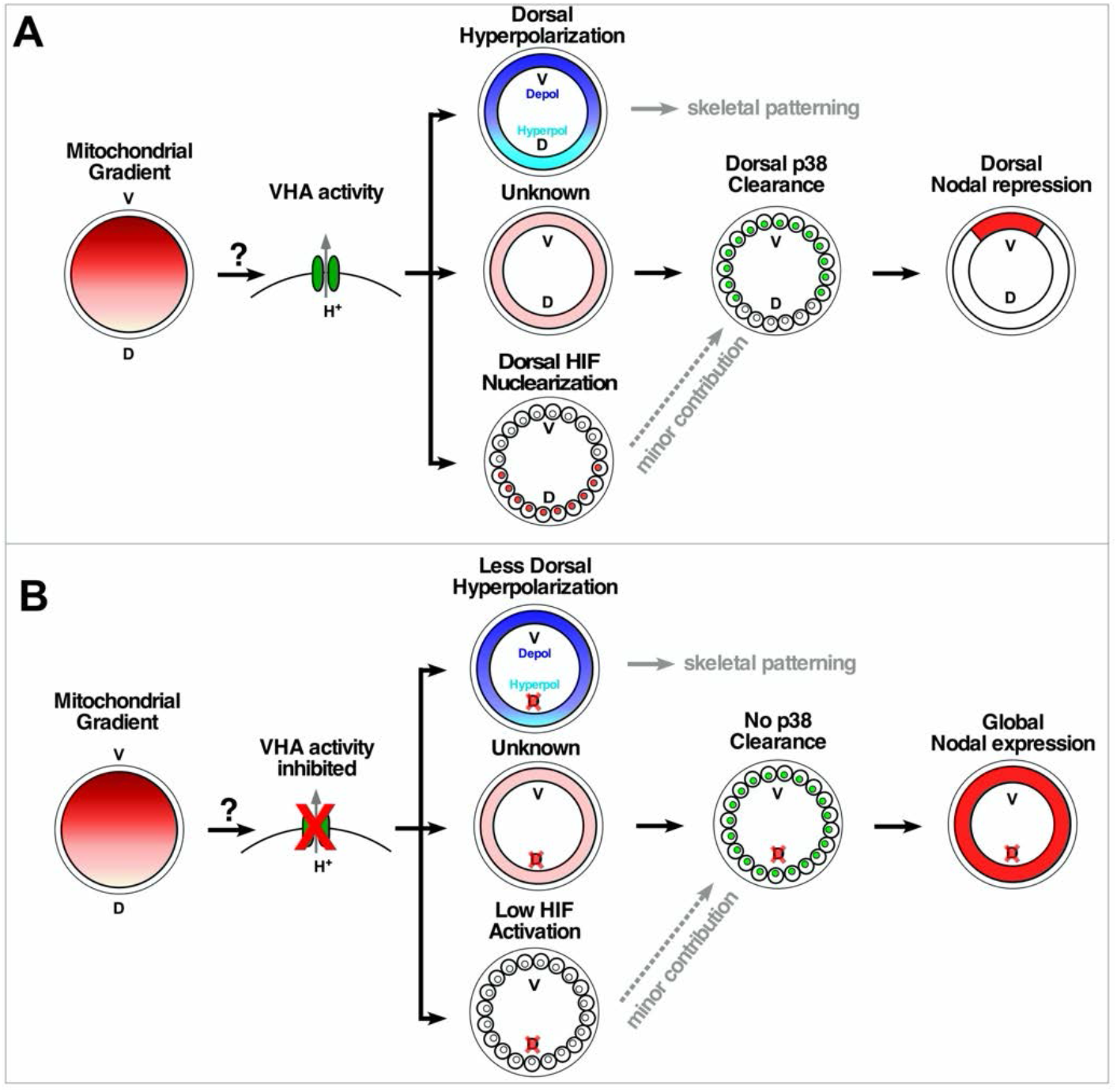
A model for VHA function in DV symmetry breaking. The model depicts control (A) and VHA inhibition (B), showing that VHA inhibition leads to global p38 activity and Nodal expression. Our results show three distinct downstream effects of the VHA in the presumptive dorsal region. Surprisingly, neither hyperpolarization nor HIF, alone or together, are required to specify dorsal, defining the necessity for an unknown third VHA-dependent component that functions to break DV symmetry upstream from p38 MAPK.

We find that VHA activity is required for the transient inhibition of p38 in the presumptive dorsal territory during DV specification, which implies that maintaining global p38 activity promotes the global onset of Nodal expression. While the requirement of p38 MAPK for Nodal expression has been previously established (Bradham and McClay, 2006b; Bradham *et al*., 2009; Cavalieri and Spinelli, 2014; Molina *et al*., 2017), these are the first data that correlate ectopic p38 activity with ectopic Nodal expression, suggesting that p38 MAPK activity is sufficient for Nodal expression. This result with VHA inhibition is corroborated by our results with hyperosmolarity, which also show a correlation of ectopic p38 activation with global Nodal expression and DV perturbation. These findings further suggest that the unknown temporal cofactor that is required for Nodal expression in addition to the spatial control mediated by p38 MAPK (Bradham and McClay, 2006b) is globally available and therefore is not also a spatial regulator.

We find that VHA activity is required for the asymmetric initiation of Nodal expression, but not to maintain Nodal asymmetry. This conclusion is based on the complete rescue of VHA-inhibited embryos by experimentally enforced asymmetric Nodal signaling, which is sufficient to rescue specification of the entire DV axis, despite the continuing presence of VHA inhibitor. Thus, regulation of the spatial extent of the ventral Nodal signaling domain does not require ongoing VHA activity, and is maintained presumably through the action of a Turing reaction-diffusion system between Nodal and its inhibitor Lefty (Duboc *et al*., 2008; Muller *et al*., 2012). Later, BMP2/4 signaling specifies the dorsal ectoderm and also acts in opposition to Nodal signaling (Duboc *et al*., 2004; Bradham *et al*., 2009; Yaguchi *et al*., 2010). These processes involving Lefty and BMP2/4 evidently proceed normally under continuing VHA inhibition as long as Nodal expression is initiated in an asymmetric manner.

Our results show that while VHA activity is required for both dorsal hyperpolarization and HIF activation, neither of these effects can account for VHA-mediated dorsal symmetry breaking. The V_mem_ gradient along the DV axis is instead required for normal skeletal patterning, while HIF makes only minor contributions to dorsal specification and skeletal patterning. Thus, the signaling events that drive DV specification downstream from the onset of Nodal expression are unaffected by differing bioelectrical states, as shown by both the rescue experiments and the potassium gluconate experiments. The lack of a functional role for the V_mem_ gradient in VHA-dependent DV specification is unexpected and surprising, since VHA-dependent biological voltage gradients are required to direct specification of the left-right axis in frogs, chickens, and zebrafish (Adams et al., 2006) and for regeneration in tadpole tails (Adams *et al*., 2007). However, in addition to its electrogenic effects, VHA is also known to impinge on numerous signaling pathways, including AMPK, PKA, Wnt/ß-catenin, Wnt/PCP, Notch, Hedgehog, TGF-ß, and mTOR (Liegeois et al., 2006; Forgac, 2007; Yan et al., 2009; Dechant et al., 2010; Hermle et al., 2010; Cotter et al., 2015; Sun-Wada and Wada, 2015; Colacurcio and Nixon, 2016; Jung et al., 2018; McGuire and Forgac, 2018), providing a wealth of alternative possibilities to pursue in future work.

The maternal mitochondrial asymmetry along the presumptive DV axis produces a redox gradient in the embryo, such that the mitochondrially-enriched, presumptive ventral territory is more oxidized (Coffman and Davidson, 2001; Coffman *et al*., 2004; Coffman *et al*., 2009; Modell and Bradham, 2011). Redox manipulation including hypoxia perturbs DV specification in Sp embryos (Czihak, 1962; 1963; Coffman and Davidson, 2001; Coffman *et al*., 2004; Coffman *et al*., 2009; Coluccio et al., 2011; Coffman et al., 2014); however, the mechanistic link between the redox gradient and Nodal has remained unclear. The VHA has been described as a redox sensor (Seidel et al., 2012), making it an attractive candidate for mediating the symmetry-breaking connection between active mitochondria and p38 MAPK/Nodal by linking redox activity to p38 MAPK regulation for spatial symmetry breaking. Our experiments position VHA activity upstream from p38 MAPK, and VHA regulation is plausibly downstream from the maternally organized redox gradient (Seidel *et al*., 2012), consistent with this model. In the future, it will be important to understand how the activity of VHA is spatially regulated during development, and to understand the mechanism by which the VHA impacts p38 MAPK activity. Finally, it will be of interest to learn how this pathway evolved and the extent of its evolutionary generalizability among echinoderms, basal deuterostomes, and anamniote vertebrates.

## Materials and Methods

### Animals, embryo cultures, and microinjection

*Lytechinus variegatus* (Lv) adults were obtained from Duke University Marine Labs (Beaufort, North Carolina) or Reeftopia (Miami, Florida). *Paracentrotus lividus* adults were obtained and used at the Observatoire Océanologique de Villefranche sur Mer, France. Spawning, microinjections, and cultures were performed by standard methods. All Lv embryos were cultured at 21°C.

### Chemical Treatments

Concanamycin A (Sigma), Bafilomycin A1 (Santa Cruz), EGA (Millipore), BEL (Sigma), Dynasore Hydrate (Sigma), SB203580 (Sigma), SB431542 (Sigma), and PT-2385 (SelleckChem or MedChem Express) were resuspended in anhydrous DMSO and diluted in artificial sea water. Suspended drugs were stored at −20°C in single use aliquots. All drug-treated embryos were compared to DMSO-treated controls; all doses were empirically optimized via dose-response experiments. The minimum effective dose for VHA inhibitors fluctuated slightly depending on the batch of drug and population of embryos; unless otherwise indicated, the dose used was about 7 nM ConA or 225 nM BafA1. These optimized concentrations are consistent with the relative IC_50_ values for these inhibitors, with ConA being approximately 20-fold more effective than BafA1 (Drose et al., 1993). SB203580 was used at 20 μM within 30 days of resuspension (Bradham and McClay, 2006b; Piacentino *et al*., 2016), and SB431542 at 5 μM (Piacentino *et al*., 2015). BEL was used at the empirically determined maximum non-lethal dose, while EGA was used at the maximum dose possible due to its poor solubility in sea water. Potassium gluconate (KGluc) was used at 30 mM. Since this dose resulted in a significant block to biomineralization, KGluc-treated embryos were washed after early gastrula stage, after DV specification is complete (Hardin *et al*., 1992; Bradham and McClay, 2006b), to allow skeletal development for morphological evaluation. PT-2385 was used at 150 μM. Embryos were treated within one hour of fertilization unless otherwise indicated. In the case of microinjected embryos, drug treatment began no more than 30 minutes after the injections were completed. For timed treatments beyond 7 hpf, timepoints were defined by morphological milestones (e.g., hatching) rather than at specific temporal intervals since the precise timing of these later milestones is slightly variable between cultures. For treatments in which embryos were fixed prior to the appearance of the anticipated phenotype, a fraction of the culture was reserved for further development to confirm the effectiveness of the treatment and the optimal dose.

### Microinjection

RNA for microinjections was prepared from template inserts in the pCS2 vector with the mMessage mMachine SP6 KIT (Ambion) and diluted in glycerol with a dextran-conjugated lineage tracer (ThermoFisher); RNA doses were empirically optimized. LvNodal, PlNodal, Lvp38-GFP, and SpHIF-mCherry were previously described (Duboc *et al*., 2004; Bradham and McClay, 2006b; Chang *et al*., 2017). Morpholinos (MOs) were purchased from Gene Tools; LvNodal MO was previously described (Bradham and McClay, 2006b). Microinjections were performed using standard approaches using a Zeiss Axiovert microscope equipped with a Narishige micromanipulator and a Sutter Instruments picospritzer.

### Immunofluorescent labeling and fluorescent in situ hybridization

Immunofluorescent labeling was performed as previously described (Bradham *et al*., 2009) using primary antibodies 295 for the ciliary band, a kind gift from Dave McClay (Duke University), 1e11 (anti-synaptotagmin B, Developmental Studies Hybridoma Bank), and anti-serotonin (Sigma). Cy2- and Cy3-conjugated secondary antibodies (Jackson Laboratories) were used at 1:300. Fluorescent in situ hybridization was carried out by standard methods. Probes for LvNodal, LvChordin, LvBMP2/4, LvTbx2/3 and LvLefty were previously described (Gross, 2003; Bradham and McClay, 2006b; Bradham *et al*., 2009; Schatzberg et al., 2015). Partial-length probes for LvIrxA, LvAtp6v1e, and LvAtp6v1h were amplified from late-gastrula stage Lv cDNA. RNA Probes were labeled with DIG (New England Biolabs) or DNP-11-UTP (Perkin Elmer) and visualized with the Tyramide Signal Amplification Kit (Perkin Elmer).

### Reporter dyes

Mitotracker Green FM, Tetramethylrhodamine (TMRM, red), Bis-(1,3-dibutylbarbituric acid)trimethine oxonol (DiBAC_4_(3), green), and Bis-(1,3-diethylthiobarbituric acid)trimethine oxonol (DiSBAC_2_(3), red) (ThermoFisher), FITC-dextran (green) (ThermoFisher) and rhodamine-dextran (red) (ThermoFisher) were prepared according to the manufacturer’s instructions. Live embryos were soaked in mitochondrial or V_mem_ indicator dyes for up to 1 hour at room temperature, then imaged.

### Microscopy

Embryos were photographed for morphology on a Zeiss Axioplan microscope at 200x magnification using either DIC optics or plane-polarized light to visualize skeletal birefringence. Skeletal images are presented as montages of several focal planes to display the entire three-dimensional skeleton in focus. Labeled FISH embryos were imaged on a Zeiss Axioplan microscope at 200x magnification using epifluorescence. Confocal microscopy was performed using an Olympus Fv10i laser scanning confocal at 600x. The Olympus software or Fiji was used to create z-stack projections for each embryo. p38-GFP (Bradham and McClay, 2006b) was imaged in live embryos with confocal microscopy every 10 minutes for 4-5 hour intervals. p38 nuclear clearance was defined by a region of more than 3 neighboring nuclei becoming indistinguishable from the background fluorescence for at least two consecutive temporal frames (>20 minutes). Reporter dyes for mitochondria and V_mem_ were detected using an Olympus DSU spinning disk confocal. Single focal planes were captured in the approximate vertical center of each embryo. To quantify the results, the signal around the circumference of the embryo was measured using manual regions of interest (ROIs) in Fiji to quantify the signal per unit area, along with measurements of the background signal collected from the corners of the image where no tissue was present. For voltage measurements, mitochondrial asymmetry was used to orient the DV axis, then measurements were made across this axis (see Fig. 5C1). For each embryo, the maximum cross-embryo net signal difference was determined after subtraction of the background as the average from each corner of the image. Each treatment group represents 10-15 embryos, and each experiment was repeated at least twice with biologically independent samples.

### Molecular constructs

A sequence encoding a constitutively active form of the Nodal receptor, LvAlk4/5/7 was generated by mutating the full-length LvAlk4/5/7 sequence (Piacentino *et al*., 2015) to change residue 271 from glutamine to aspartic acid (Lapraz et al., 2015) using standard molecular techniques, resulting in LvAlk4/5/7 Q271D (AlkQD), then cloned into pCS2 vector for *in vitro* transcription.

### qPCR

Total RNA was collected using the RNEasy Mini kit (QIAGEN) and DNAse-treated with the RNAse-free DNAse Set (QIAGEN). Samples were reverse transcribed using the TaqMan Reverse Transcription Kit (ThermoFisher); qPCR was performed using SYBR Green PCR Master Mix (ThermoFisher) in an ABI 7900ht qPCR thermocycler. Each data point represents a minimum of two biologically independent replicates (each in triplicate, for a minimum of 6 measurements). ΔC_T_ values were normalized to LvSetmar expression (Hogan *et al*., 2020); ΔΔC_T_ values were determined by comparison to controls. Primer sequences used were previously reported (Gildor and Ben-Tabou de-Leon, 2015; Schatzberg *et al*., 2015), except those for LvAtp6v0a1 (forward: 5’-GCTCCTGCTCCTAGGGAGAT-3’ and reverse: 5’-TCTCCATCGTCCATCTCATC-3’).

## Acknowledgements

We thank Dave McClay (Duke University) for the gift of antibodies and for helpful comments on the manuscript, Jenifer Croce and Guy Lhomond (Observatoire Océanologique de Villefranche sur Mer, France) for access to *P. lividus* sea urchins, and Mike Levin (Tufts University) for helpful advice. This work was supported by NSF IOS grants 1257825 and 1656752 (CAB). DS was partially supported by Warren-McCloud and Terner awards (Biology Department, Boston University); PR, SEH, VS, and SK were partially supported by undergraduate research opportunities program (UROP) awards (Boston University).

## Author Contributions

The project was conceived by CAB. The experiments were performed by DS, CFT, PR, SEH, VS, MLL, BD, SK, and DTZ. The data were analyzed by DS, CFT, SEH, VS, LD, and CAB. The manuscript was written by DS and CAB and edited by all co-authors.

