## Supplemental Figures & Table for "V-type H^+^ ATPase Activity is Required for Embryonic Dorsal-Ventral Symmetry Breaking"

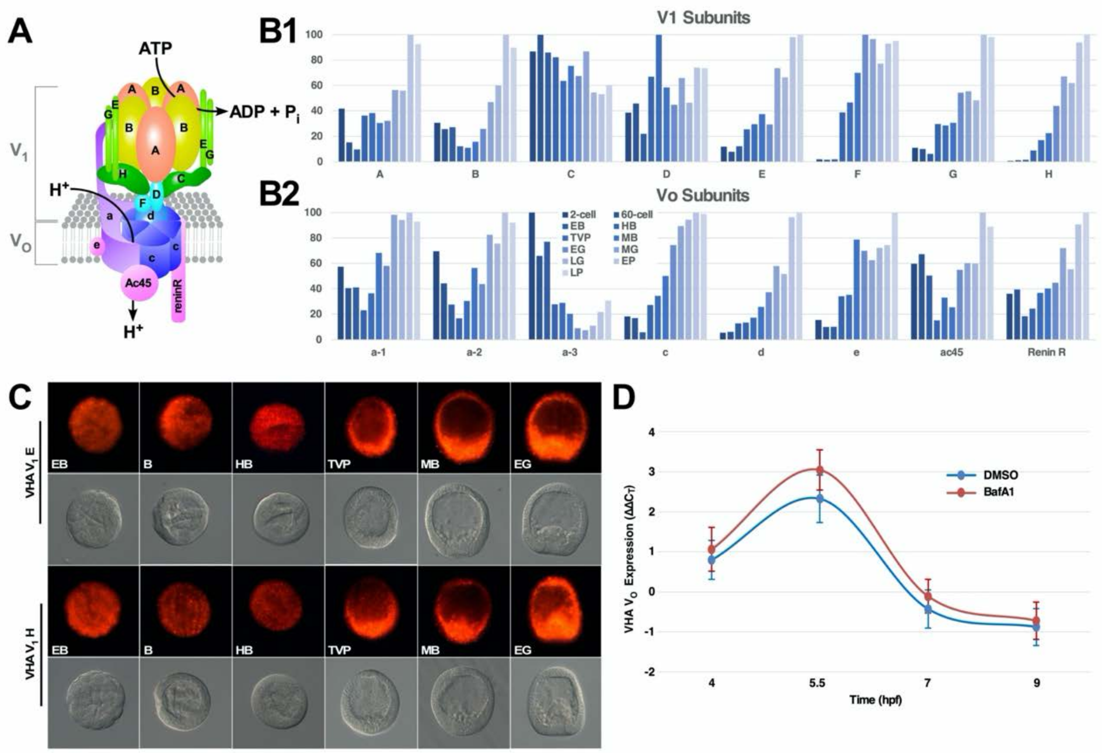


**Figure S1. All VHA subunits are highly expressed after EB stage (4 hpf) in Lv embryos.**

**A.** The schematic illustrates the V-ATPase $V_O$ and $V_1$ domains, and the subunits that comprise each domain (image adapted from Sun-Wada and Wada, 2015).
**B.** The temporal expression of the subunits that compose the $V_1$ (B1) or $V_O$ (B2) subassemblies in Lv embryos is shown as their percent maximum expression. All subunits exhibit non-zero expression at all timepoints, while expression is strongly increased for subunits F, H, and c after 4 hpf (EB stage).
**C.** Fluorescent in situ hybridization (FISH) for VHA $V_1$ subunits E and H is shown along with the associated DIC images at the indicated developmental stages.
**D.** The time course for expression of VHA $V_O$ subunit a-1 is shown for DMSO-treated controls (blue) and Baf-treated embryos (red) at the indicated time points spanning EB (early blastula, 4 hpf), HB (hatched blastula, 7 hpf) and TVP (thickened vegetal plate, 9 hpf) as the average $\Delta\Delta C_T$ ± sem compared to 4 hpf. The results show that BafA1 treatment does not affect VHA $V_O$ subunit a-1 expression. See Fig. 2 for other developmental stage abbreviations.

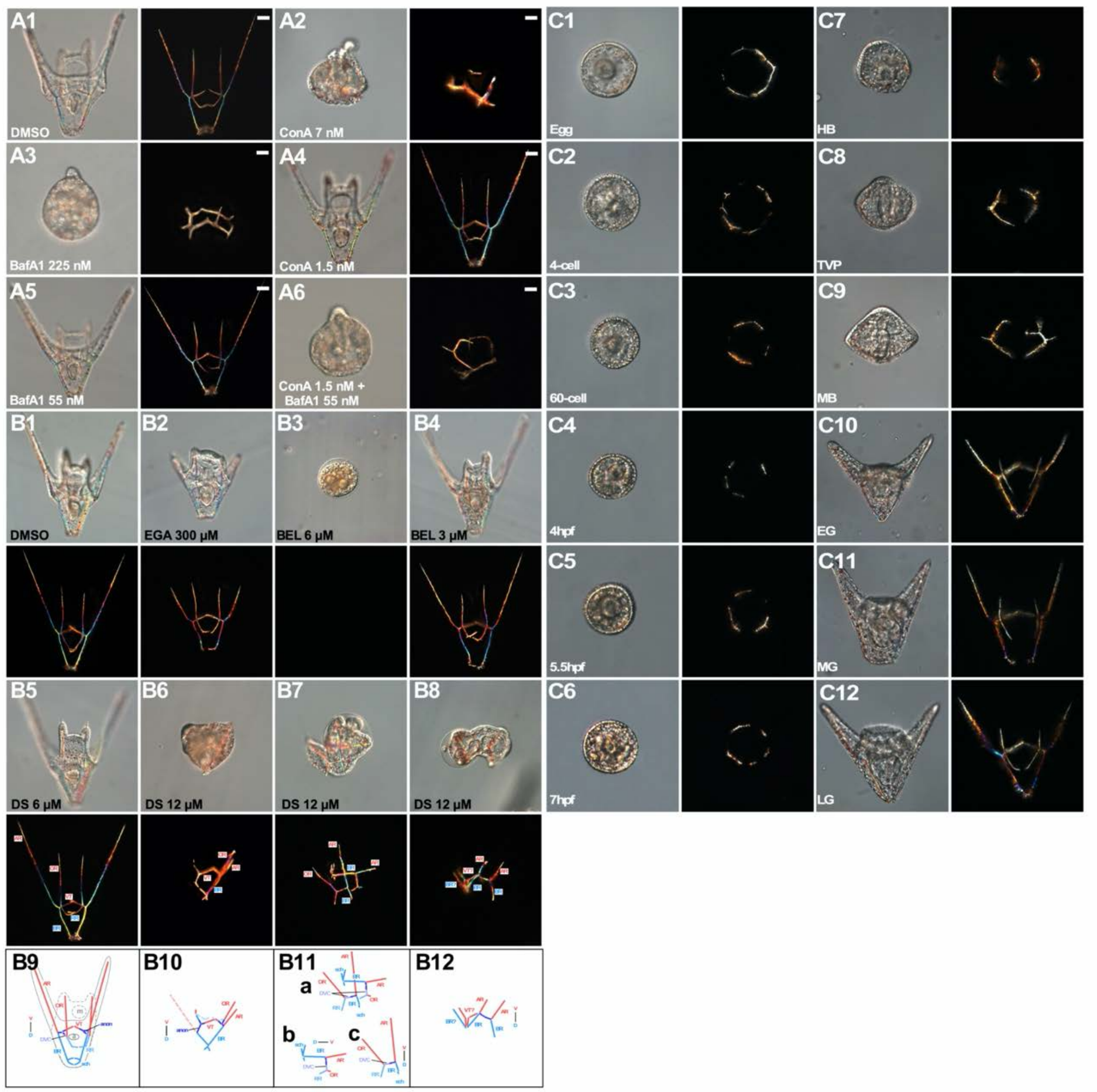


**Figure S2. VHA inhibitors ConA and BafA1 have additive effects when combined at suboptimal doses, are not phenocopied by endocytosis inhibitors, and become ineffective when added after hatched blastula stage**

**A.** Embryos were treated as indicated with optimal (A2-3) and suboptimal (A4-6) doses of VHA inhibitors, then imaged at 48 hpf; DIC (right) and skeletal birefringence montage images (left) are shown.

**B.** Embryos were treated with the indicated endocytosis inhibitors at the indicated doses, then imaged at 48 hpf, with DIC (upper panels) and skeletal birefringence montage images (lower panels) shown (B1-8). For DS-treated embryos, skeletal patterns are also schematized, with dorsal (blue) and ventral (red) skeletal elements indicated (B9-12), along with initial triradiates (dark blue) and the DVCs that are perpendicular to the DV axis depicted in violet. The exemplar shown in B7 and B11 exhibits perpendicularly oriented skeletal halves, which are depicted schematically together (a) and individually (b-c) for better clarity.

**C.** Embryos were treated with ConA at the indicated time points, then imaged at 48 hpf; DIC (1) and skeletal birefringence montage images (2) are shown. Bilateral skeletons are evident beginning at HB (G). See Figure 2 for quantification and developmental stage abbreviations. All larvae are oriented with ventral upward when the DV axis is discernable.

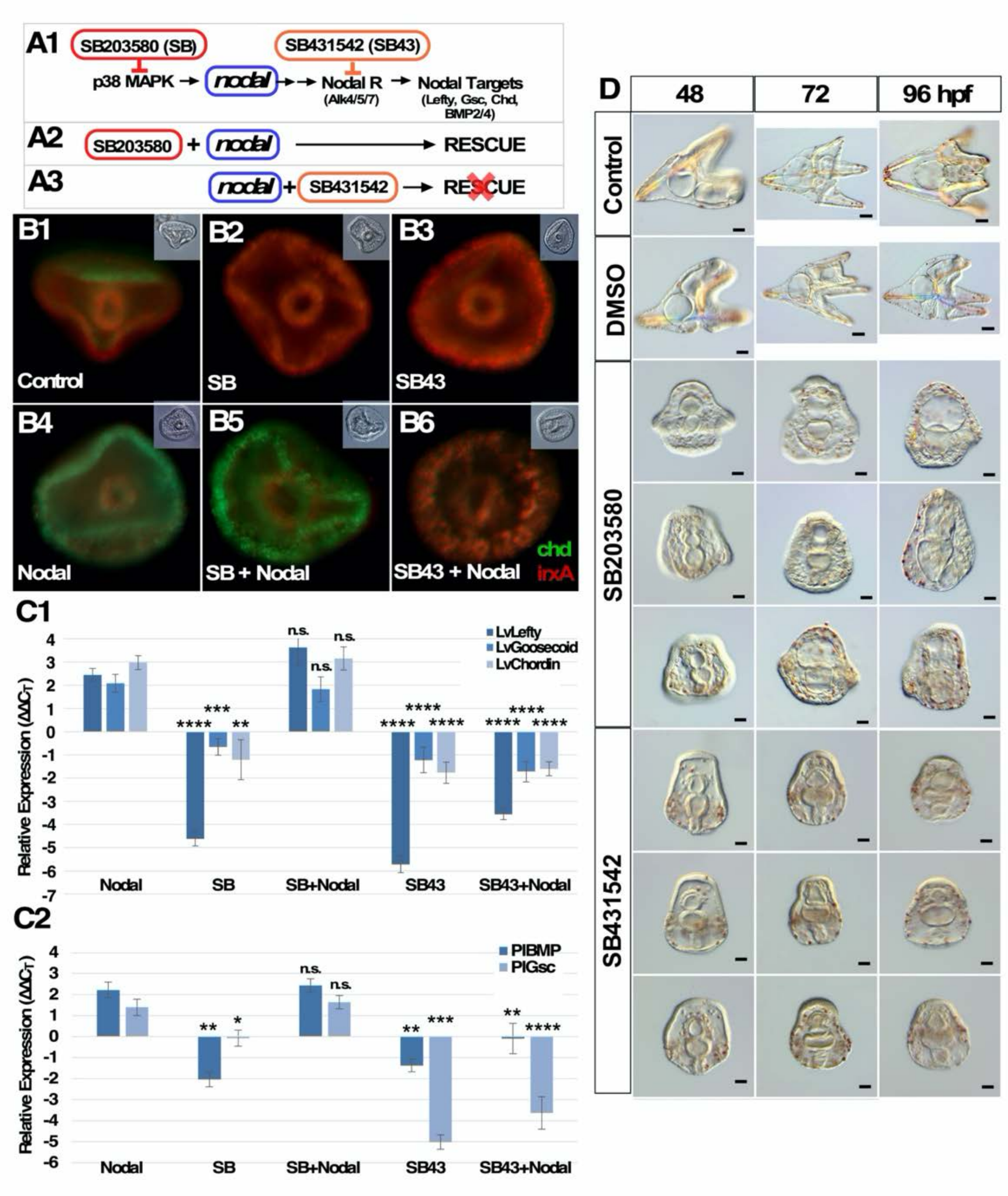


**Figure S3. The p38 MAPK inhibitor SB203580 acts upstream from Nodal expression and results in radialized embryos in *Paracentrotus lividus*.**

**A.** The ventral specification pathway in Lv is shown, with the expected targets of p38 MAPK and Nodal receptor Alk4/5/7 inhibitors indicated (A1). SB203580 inhibits p38 MAPK upstream from Nodal transcription; thus, Nodal overexpression should rescue SB203580-treated embryos (A2). In contrast, SB431542 inhibits the Nodal receptor Alk4/5/7 downstream from the action of Nodal ligand and therefore should not be rescued by Nodal overexpression (A3).

**B.** *L. variegatus* embryos were treated with the p38 MAPK inhibitor SB203580 (SB) or the Alk4/5/7 inhibitor SB431542 (SB43), with or without Nodal overexpression as indicated, then analyzed at 18 hpf for spatial gene expression of ventral chordin (green) and dorsal irxA (red) using FISH; corresponding DIC images are inset. Treatment with SB or SB43 alone results in global expression of the dorsal marker (B2, B3), while Nodal overexpression alone results in global expression of the ventral marker (B4). Nodal rescues ventral gene expression when combined with SB (B5) but not combined with SB43 (B6), consistent with SB203580 inhibition functioning upstream and not downstream from Nodal expression (A).

**C.** *L. variegatus* (C1) or *P. lividus* (C2) **e**mbryos were treated as in B, then analyzed for expression of the indicated Nodal target genes at 18 hpf via qPCR. The data are shown as the average $\Delta\Delta C_T$ compared to DMSO-treated control embryos ± s.e.m. Statistical significance compared to Nodal overexpression was analyzed with paired student t tests; * $p < 0.05$, ** $p < 0.005$, *** $p < 0.0005$, **** $p < 0.000001$. These results show that p38 MAPK inhibition is rescued by Nodal overexpression, while inhibition of Nodal's receptor Alk4/5/7 is not rescued by Nodal overexpression, as schematized in A. These results along with B. are inconsistent with a model in which SB203580 inhibits Alk4/5/7.

**D.** Control, DMSO- and drug-treated *P. lividus* exemplar embryos are shown at 48, 72, and 96 hpf as indicated; representative morphologies are depicted. Phenotypes with SB20350 that are similar to those described by Molina et al. (Molina et al., 2017) were not observed. See also Fig. 4.

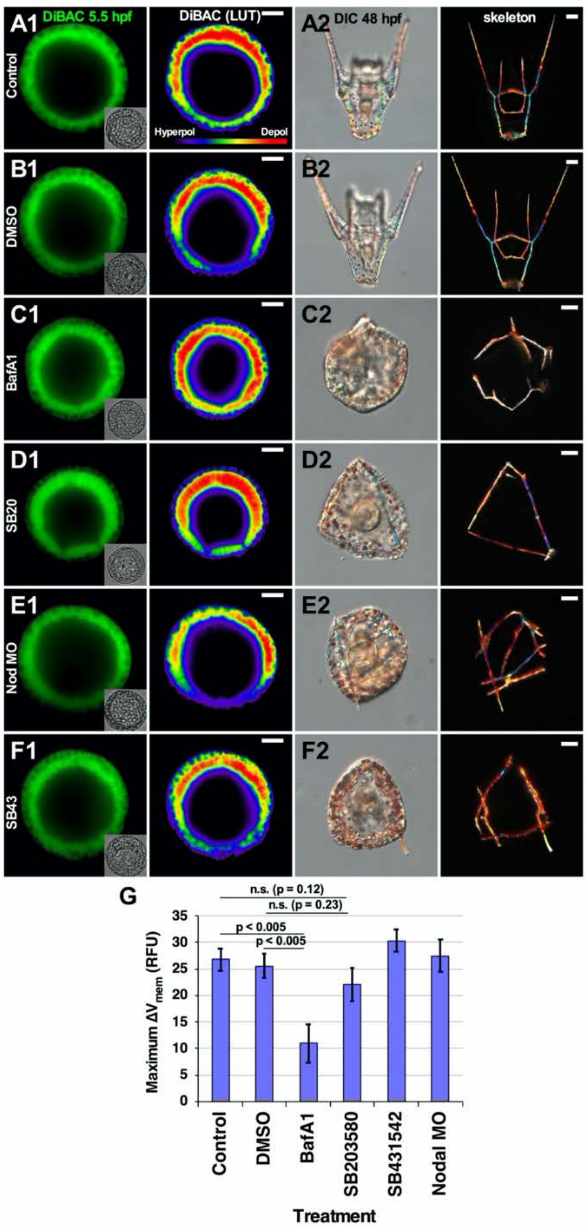


**Figure S4. The endogenous $V_{mem}$ gradient along the DV axis does not require p38-MAPK activity, Nodal expression, or Nodal signaling.**

Control (A), DMSO- (B), BafA1-(C), SB203580- (D), SB431542-treated (F) and Nodal MO-injected (E) embryos were analyzed for $V_{mem}$ using DiBAC at 7 hpf and shown as raw fluorescence (1, left) or with pseudocoloring (2, right), and for morphology at 48 hpf, with DIC (2, left) and skeletal birefringence montages (2, right) shown. The maximum cross-embryo $V_{mem}$ signal was quantified as the average ± s.e.m.; $n \geq 12$ embryos per condition; p values were determined using paired student t tests in comparison with control or DMSO (G). See also Fig. 5.

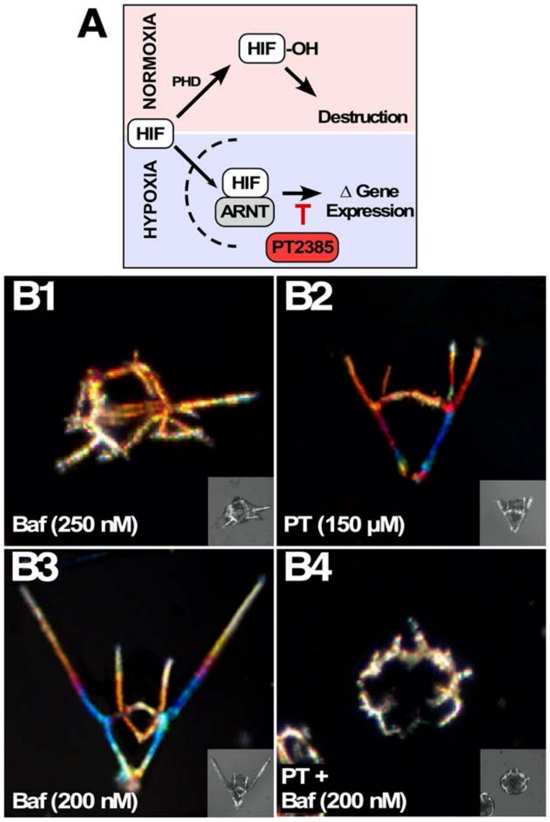


**Figure S5. HIF inhibition, while functionally additive to suboptimal VHA inhibition, is insufficient alone to perturb the DV axis.**

**A.** HIF is hydroxylated by proline-directed hydroxylases (PHD) under normoxia, promoting HIF degradation. Under hypoxia, PHD proteins are inhibited, and HIF translocates to the nucleus and dimerizes with ARNT to modulate transcription. PT-2385 is a specific inhibitor of HIF activity.

**B.** HIF is not required for DV specification (B2), but HIF inhibition has additive effects with suboptimal doses of BafA1 (B3-4). Embryos treated with the indicated doses of BafA1 and/or PT-2385 are shown at 48 hpf as skeletal birefringence montage images with the corresponding DIC images inset. See also Fig. 5.

**Table S1. VHA subunit genes**

| Domain | Subunit | SPU | Name | Alt. Name |
|---|---|---|---|---|
| $V_1$ | A | SPU_016882 | LvAtp6v1a2 | |
| $V_1$ | B | SPU_006017, SPU_016414 | LvAtp6v1b2 | LvVha55 |
| $V_1$ | C | SPU_006308 | LvAtp6v1c1 | |
| $V_1$ | D | SPU_012628 | LvAtp6v1d | |
| $V_1$ | E | SPU_009588 | LvAtp6v1e | LvVha26 |
| $V_1$ | F | SPU_012784 | LvAtp6v1f | |
| $V_1$ | G | SPU_002356 | LvAtp6v1g3 | |
| $V_1$ | H | SPU_000364 | LvAtp6v1h | |
| $V_O$ | a-1 | SPU_001860 | LvAtp6v0a1 | |
| $V_O$ | a-2 | SPU_002051 | LvAtp6v0a1_1 | |
| $V_O$ | a-3 | SPU_023764, SPU_020497 | LvAtp6v0a1_3 | |
| $V_O$ | c | SPU_016993 | LvAtp6v0c | LvAtp6v0b; 21 kDa proteolipid |
| $V_O$ | d | SPU_013772 | LvAtp6v0d | |
| $V_O$ | e | SPU_018565 | LvAtp7v0e | LvVhaac39 |
| $V_O$ | ac45 | SPU_012695 | LvAtp6ap1 | |
| $V_O$ | Renin R | SPU_003104 | LvAtp6ap2 | |
